# Digitally immune optimised haemagglutinin with nanocage plug-and-display elicits broadly neutralising pan-H5 influenza subtype vaccine responses

**DOI:** 10.1101/2024.11.14.623359

**Authors:** Chloe Qingzhou Huang, Rory A. Hills, George W. Carnell, Sneha Vishwanath, Ernest T. Aguinam, Andrew C.Y. Chan, Phil Palmer, Laura O’Reilly, Paul Tonks, Nigel Temperton, Simon D.W. Frost, Laurence S. Tiley, Mark R. Howarth, Jonathan L. Heeney

**Affiliations:** Laboratory of Viral Zoonotics, Department of Veterinary Medicine, University of Cambridge, Madingley Road, Cambridge, CB3 0ES, UK; Department of Pharmacology, University of Cambridge, Tennis Court Road, Cambridge, CB2 1PD, UK; Department of Biochemistry, University of Oxford, South Parks Road, Oxford, OX1 3QU, UK; Viral Pseudotype Unit, Medway School of Pharmacy, The Universities of Kent and Greenwich at Medway, Central Avenue, Chatham, ME4 4TB, UK; Department of Veterinary Medicine, University of Cambridge, Madingley Road, Cambridge, CB3 0ES, UK; London School of Hygiene and Tropical Medicine, Keppel Street, London, WC1E 7HT, UK; Microsoft Health Premonition, One Microsoft Way, Redmond, WA, 98052, USA; DIOSynVax Ltd, Janelle House, 6 Hartham Lane, Hertford, SG14 1QN, UK

**Author notes:** These authors contributed equally (Chloe Qingzhou Huang, Rory A. Hills). Corresponding authors: Jonathan L. Heeney Mark R. Howarth Laurence S. Tiley.

## Abstract

The increasing global spread of the highly pathogenic avian influenza (HPAI) A/H5 viruses poses a serious public health threat. Circulating clade 2.3.4.4b viruses have demonstrated rapid transcontinental dissemination, extensive reassortment, epizootic spread and potential sustained mammal-to-mammal transmission, signifying a heightened risk of becoming a human pathogen of high consequence. A broadly protective, future-proof vaccine against multiple clades of H5 influenza is urgently needed for pandemic preparedness. Here, we combine two novel vaccine technologies to generate a Digitally Immune Optimised and Selected H5 antigen (DIOSvax-H5_inter_) displayed multivalently on the mi3 nanocage using the SpyTag003/SpyCatcher003 conjugation system. Mice immunised with DIOSvax-H5_inter_ Homotypic Nanocages at low doses demonstrate potent, cross-clade neutralising antibody and T cell responses against diverse H5 strains. DIOSvax-H5_inter_ Homotypic Nanocages provide a scalable vaccine candidate with the potential for pan-H5 protection against drifted or newly emergent H5 strains. This World Health Organization preferred product characteristic is essential for prospective strategic stockpiling in the pre-pandemic phase.

## INTRODUCTION

Highly pathogenic avian influenza (HPAI) viruses are characterised by the acquisition of a polybasic cleavage site in their surface glycoprotein haemagglutinin (HA)^1,2^. Recognition and cleavage by ubiquitous cellular proteases permit productive infection beyond the respiratory tract, facilitating systemic infection and increased viral pathogenicity^1,2^. HPAI viruses of the H5 subtype of the A/goose/Guangdong/1/1996 (gs/GD) lineage originated in domestic poultry in southeast Asia^2^. Evolution of the HA gene and multiple reassortment with low pathogenicity avian influenza viruses (LPAI) enabled adaptation for transmission of clade 2 viruses to wild aquatic birds that expanded the geographic range of the virus within the continent^3^. Since 2018, re-emergence of a novel reassortant of clade 2.3.4.4b in migratory bird populations with enhanced genetic plasticity and host adaptation has led to a global panzootic with infections in over 150 avian and 50 mammalian species^4^. Potential sustained mammal-to-mammal transmission and human infections from marine and terrestrial mammals, including food producing dairy cattle, highlight the urgent need for pandemic preparedness^4^.

A critical task of pandemic preparation is stockpiling effective vaccines^5^. Based on global surveillance of circulating HPAI A/H5 isolates, the World Health Organization provides an annually updated list of representative candidate vaccine virus (CVV) strains^5^. Strain-specific inactivated vaccines have been traditionally stockpiled for zoonotic influenza viruses^6,7^. mRNA vaccines matched to the latest CVV H5 A/Astrakhan/3212/2020 (H5Ast20) have also been developed, which can be manufactured on a reduced timescale in response to a pandemic^8–10^. However, as recently observed for the coronavirus disease 2019 (COVID-19) pandemic, accelerated evolution of RNA viruses in an expanding host reservoir can lead to drastic antigenic changes resulting in vaccine mismatch^11^. Adaptive mutations that facilitate human-to-human transmission, such as greater HA stability and binding to human α2,6-linked sialic acids and enhanced viral polymerase function, may be anticipated, but the exact composition of mutations in a specific spillover event cannot be precisely predicted^4^. A broad H5 vaccine offering cross-clade protection would effectively abrogate the need for perfect antigenic match with a pre-emergent pandemic strain of unknown identity.

To address this problem, we combined a novel computationally designed vaccine antigen using Digitally Immune Optimised Synthetic Vaccine (DIOSynVax) antigen design technology^12–16^, with the Plug-and-Display SpyTag003/SpyCatcher003-mi3 nanoassembly platform^17,18^, to develop an optimised H5 HA nanocage vaccine. Using the DIOSynVax computational pipeline, we generated a novel, recombinant H5 immunogen with antigenic determinants representative across different clades of the H5 subtype, DIOSvax-H5_inter_^15,16^. DIOSvax-H5_inter_ was generated as a soluble protein or as a self-assembling Homotypic Nanocage and compared to the CVV H5Ast20 as a Homotypic Nanocage in mice. Upon administration of two doses, only the group immunised with DIOSvax-H5_inter_ Homotypic Nanocage elicited potent neutralisation of all 12 tested H5 clades and subclades, as well as demonstrating robust CD4^+^ T cell responses against two antigenically distant H5 clades that emerged both before or after the antigen was designed. These characteristics demonstrate that the DIOSvax-H5_inter_ Homotypic Nanocage has the potential to be employed as a broadly protective pre-pandemic vaccine for stockpiling that offers variant-proof protection against the prospective threat of an H5 avian flu pandemic in humans before it begins.

## RESULTS

### *In silico* design of pan-H5 antigen

A computationally-optimised H5 antigen, termed DIOSvax-H5_inter_, was generated through the DIOSynVax phylogeny-based computational pipeline for designing broadly reactive immunogens^13,15,16^. Nucleotide sequences of influenza HA proteins were downloaded from the Global Initiative on Sharing All Influenza Data (GISAID) EpiFlu^TM^ database^19^ (June 2018) and used as input to generate a multiple sequence alignment. The resulting alignment was used as input for phylogenetic tree reconstruction. The resulting tree was further analysed using HyPhy^20^ to reconstruct the statistically-inferred and phylogenetically-optimised DIOSvax-H5_inter_ design.

The DIOSvax-H5_inter_ polypeptide sequence differs from the circulating clade 2.3.4.4b CVV H5Ast20 strain with mutations spanning both head and stem domains (Fig. 1a). Comparing DIOSvax-H5_inter_ with wild-type viral isolates from 12 major clades and subclades of the H5 subtype shows that DIOSvax-H5_inter_ is positioned phylogenetically between clades 2.1.3.2 and 2.2 (Fig. 1b). Comparison of DIOSvax-H5_inter_ with other wild-type strains from these selected H5 clades showed the highest sequence similarity of 98.4% with the clade 2.2 A/whooper swan/Mongolia/244/2005 strain. The lowest sequence similarity of 83.9% was observed with the LPAI American non-gs/GD lineage A/chicken/Mexico/07/2007 strain (Fig. S1). There was a total of 41 residues that were distinct in DIOSvax-H5_inter_ relative to the CVV H5Ast20 (Fig. 1c).

**Fig. 1.**
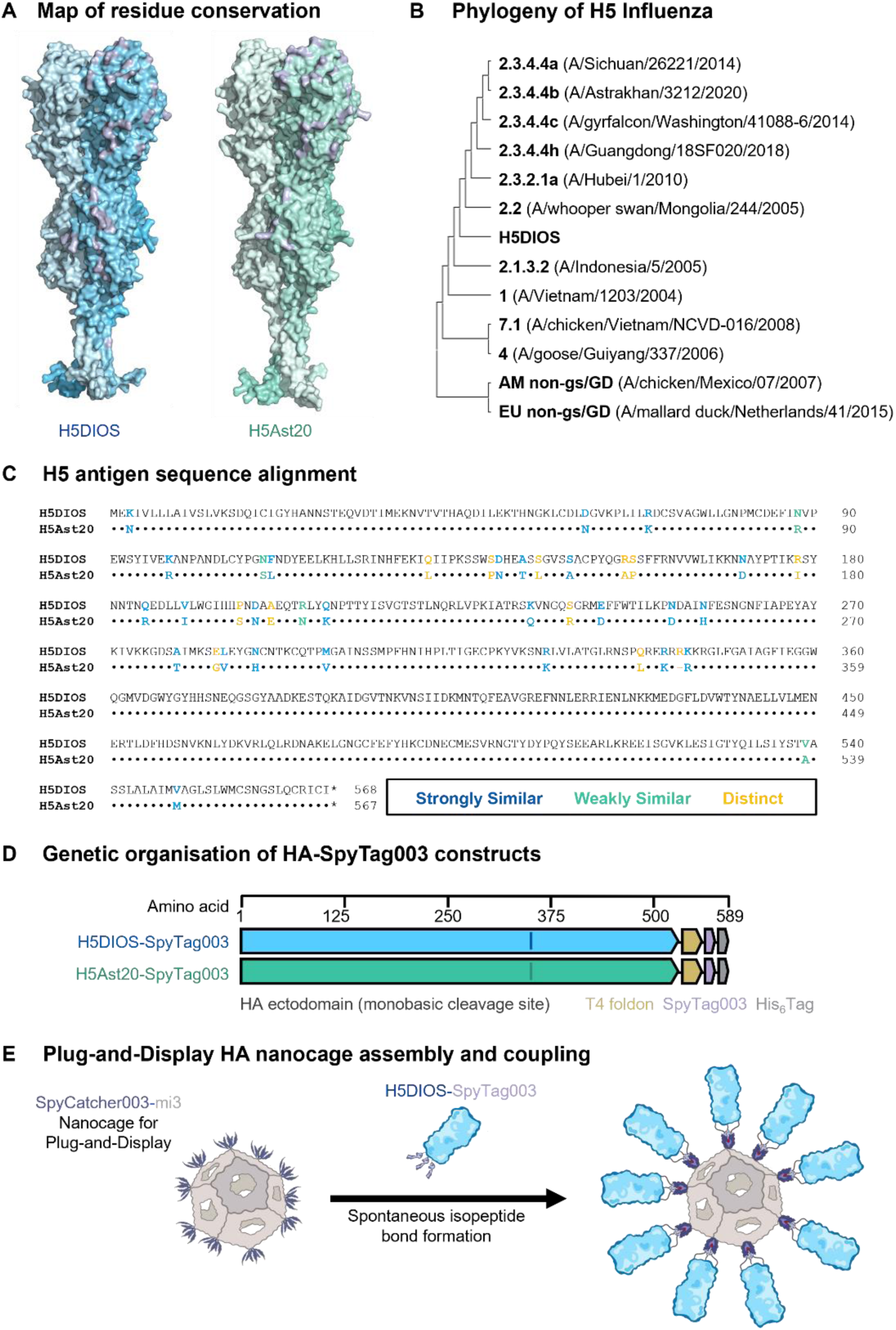
*In silico* design and assembly of H5 Nanocage vaccines. **A)** Structural models of HA antigens. Conservation of residues between full-length DIOSvax-H5_inter_ (H5DIOS) (blue) and H5Ast20 (green) mapped onto the van der Waals surface of HA homotrimers. Residue substitutions are represented in purple. Structural prediction was performed by AlphaFold 3^29^ and displayed using PyMOL. **B)** Phylogenetic tree of HA amino acid sequence identity of H5 viruses used in this study. **C)** Amino acid sequence alignment of DIOSvax-H5_inter_ and H5Ast20 by Clustal Omega v.1.2.4, numbered according to DIOSvax-H5_inter 30_. **D)** Genetic organisation of DIOSvax-H5_inter_-SpyTag003 and H5Ast20-SpyTag003 constructs, indicating the HA ectodomain, T4 foldon, tag and linker locations. **E)** Schematic of Plug-and-Display nanocage vaccine assembly of DIOSvax-H5_inter_ Homotypic Nanocage.

### Preparation and characterisation of computationally-designed H5 nanocages

mi3 is a computationally-designed, self-assembling 60-mer protein nanocage which can be expressed and purified in a scalable fashion from *Escherichia coli* and shows high stability^21,22^. Genetic fusion of mi3 to SpyCatcher003 enables high efficiency coupling through spontaneous formation of an isopeptide bond to SpyTag003-antigens^18^ of various symmetries^23^. Multivalent display presents antigens in highly repetitive arrays that recapitulate features common to viral surfaces. This arrangement enhances immune potency through mechanisms including enhanced uptake by antigen-presenting cells and efficient cross-linking by B cell receptors^24,25^. This Plug-and-Display nanocage platform has been used to develop influenza vaccines^23^ and pan-sarbecovirus vaccines with potent immunogenicity in animal models^26–28^.

We genetically fused the *in silico* optimised DIOSvax-H5_inter_ ectodomain to a T4 bacteriophage fibritin foldon^31^ flanked by two flexible glycine-serine linkers, along with SpyTag003 and His_6_ tag^32^ (Fig. 1d). Fusion to the fibritin foldon domain promoted trimerisation of the HA protein and mimicked HA’s natural conformation on virion surfaces. The characteristic polybasic cleavage site of HPAI A/H5 viruses was converted to a monobasic form through deletion of residues 341-345 (RRRKK), which stabilised HA in the pre-fusion (HA0) conformation by preventing uncontrolled cleavage by constitutive proteases present in the expression cell line^33^. We also cloned the H5Ast20 ectodomain into the same vector backbone and generated the monobasic mutant through deletion of residues 341-344 (KRRK). This HA was selected as it is the latest H5 CVV listed by the World Health Organization for stockpiling and has been used as the vaccine strain in inactivated^7^, mRNA^8–10^ and viral-vectored platforms^34,35^. Both HA constructs were efficiently expressed and secreted by mammalian Expi293F^TM^ cells and affinity-purified via the terminal His_6_ tag.

Purified HA-SpyTag003 was coupled to SpyCatcher003-mi3 (Fig. 1e) by mixing the two proteins at different molar ratios for 24 h at 4 °C. Minimal residual unconjugated HA-SpyTag003 antigens at 1:1 and 3:2 molar ratios were observed following SDS-PAGE/Coomassie (Fig. 2a and 2b), indicating that coupling is occurring predominantly through all three SpyTag003 moieties. The diffuse HA-SpyTag003 bands were consistent with the natural heterogeneity of HA glycosylation, as confirmed by digestion with Peptide N-Glycosidase F (PNGase F). Removal of N-linked glycans produced a decrease in apparent molecular weight and sharpening of the bands (Fig. 2c). Dynamic light scattering (DLS) demonstrated an increased hydrodynamic diameter for SpyCatcher003-mi3 coupled to either HA immunogen, relative to the nanocage on its own (Fig. 2d). Transmission electron microscopy (TEM) confirmed the integrity of both nanoparticles (Fig. 2e).

**Fig. 2.**
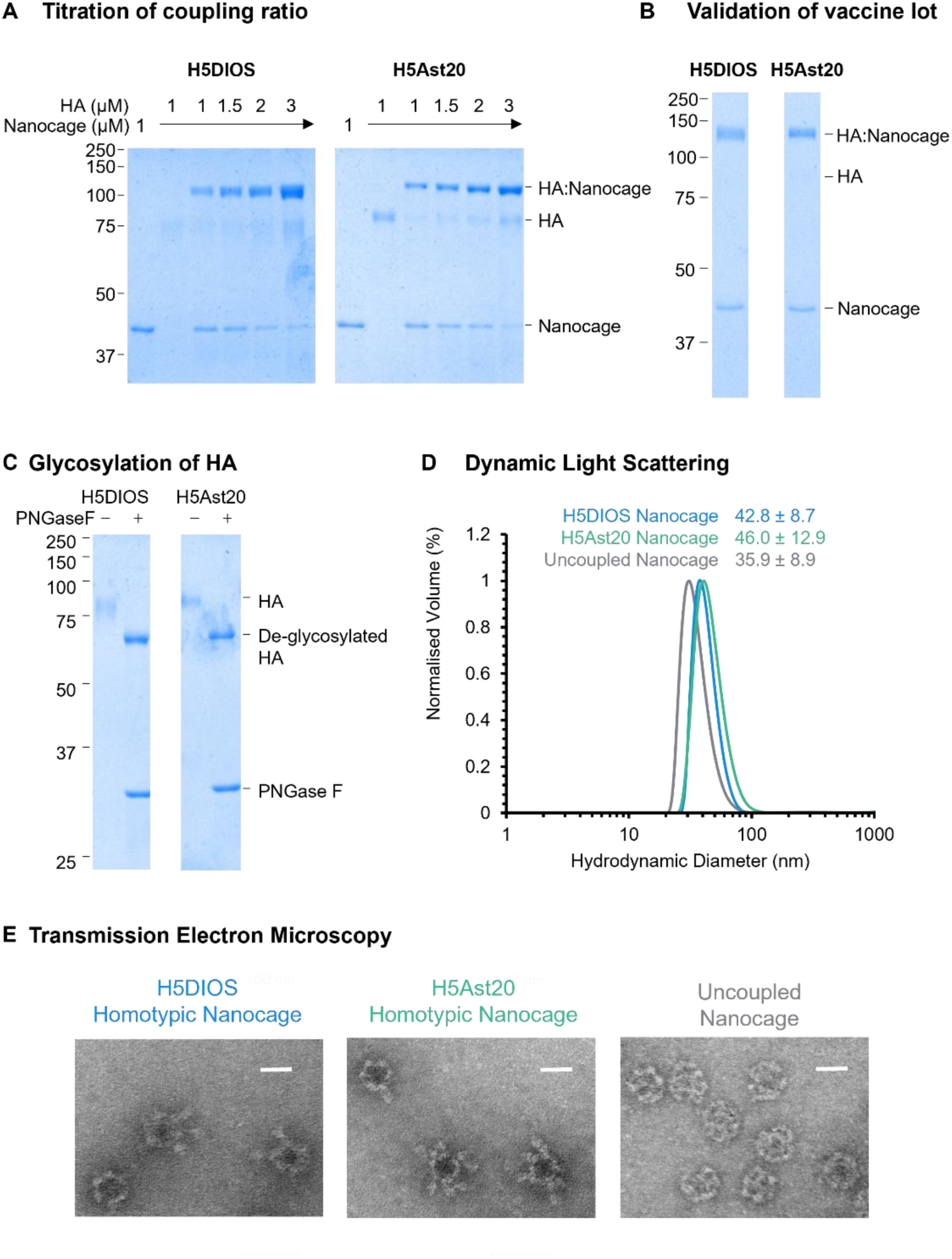
Production and characterisation of H5 Nanocage vaccines. **A)** Titration of coupling of HA-SpyTag003 to SpyCatcher003-mi3 Nanocage at different molar Nanocage:HA ratios, analysed by SDS–PAGE/Coomassie. **B)** Validation of the DIOSvax-H5_inter_ (H5DIOS) Homotypic Nanocage and H5Ast20 Homotypic Nanocage vaccine lots by SDS–PAGE/Coomassie. **C)** Analysis of HA-SpyTag003 with SDS–PAGE/Coomassie, with or without PNGase F de-glycosylation. Molecular weight markers are in kDa. **D)** Dynamic light scattering of SpyCatcher003-mi3 nanocages. The mean hydrodynamic diameter is shown ± 1 s.d.; *n* = 3. **E)** Transmission electron micrographs of negative-stained SpyCatcher003-mi3 nanocages. 50,000× magnification, scale bar = 20 nm.

To characterise the antigenicity of vaccine immunogens, we measured binding using an enzyme-linked immunosorbent assay (ELISA) to immobilised H5 soluble proteins or Homotypic Nanocages by the broadly neutralising anti-HA monoclonal antibody CR9114^36^ and by matched or closely matched mouse sera generated in-house from previous vaccination studies (Fig. S2). CR9114 is a broadly neutralising antibody targeting a conserved, 3-dimensional epitope in the HA stem of all influenza A viruses^36^. Matched or closely matched mouse sera should contain a mixture of antibodies that are head-or stem-specific, subtype-specific or broad, and neutralising or non-neutralising. Binding to CR9114 and matched or closely matched sera was observed for all three immunogens, which indicated that the HA proteins were correctly folded and multiple epitopes were exposed and accessible for binding. We noticed a substantial decrease in the maximal OD_450_ and IgG titre area under curve (AUC) for the DIOSvax-H5_inter_ antigen when expressed on Homotypic Nanocages, likely as a result of reduced accessibility by binding antibodies during multivalent display.

### Computationally-designed H5 nanocages elicit TNF^+^IFNγ^−^CD4^+^ T cell responses against multiple H5 clades

We compared the immunogenicity of DIOSvax-H5_inter_ and H5Ast20 Homotypic Nanocages, as referenced against DIOSvax-H5_inter_ Soluble protein. Doses were normalised by the number of HA trimer molecules, to allow comparison between soluble and multimerised antigens and facilitate an equimolar amount of SpyCatcher003-mi3 nanocages with similar occupancy for both Homotypic Nanocages. Groups of six mice were immunised intramuscularly at weeks 0 and 4 with 0.5 μg of HA-SpyTag003 antigen adjuvanted 1:1 with AddaVax. Terminal bleed and splenocyte harvest were performed at week 8 (Fig. 3a).

**Fig. 3.**
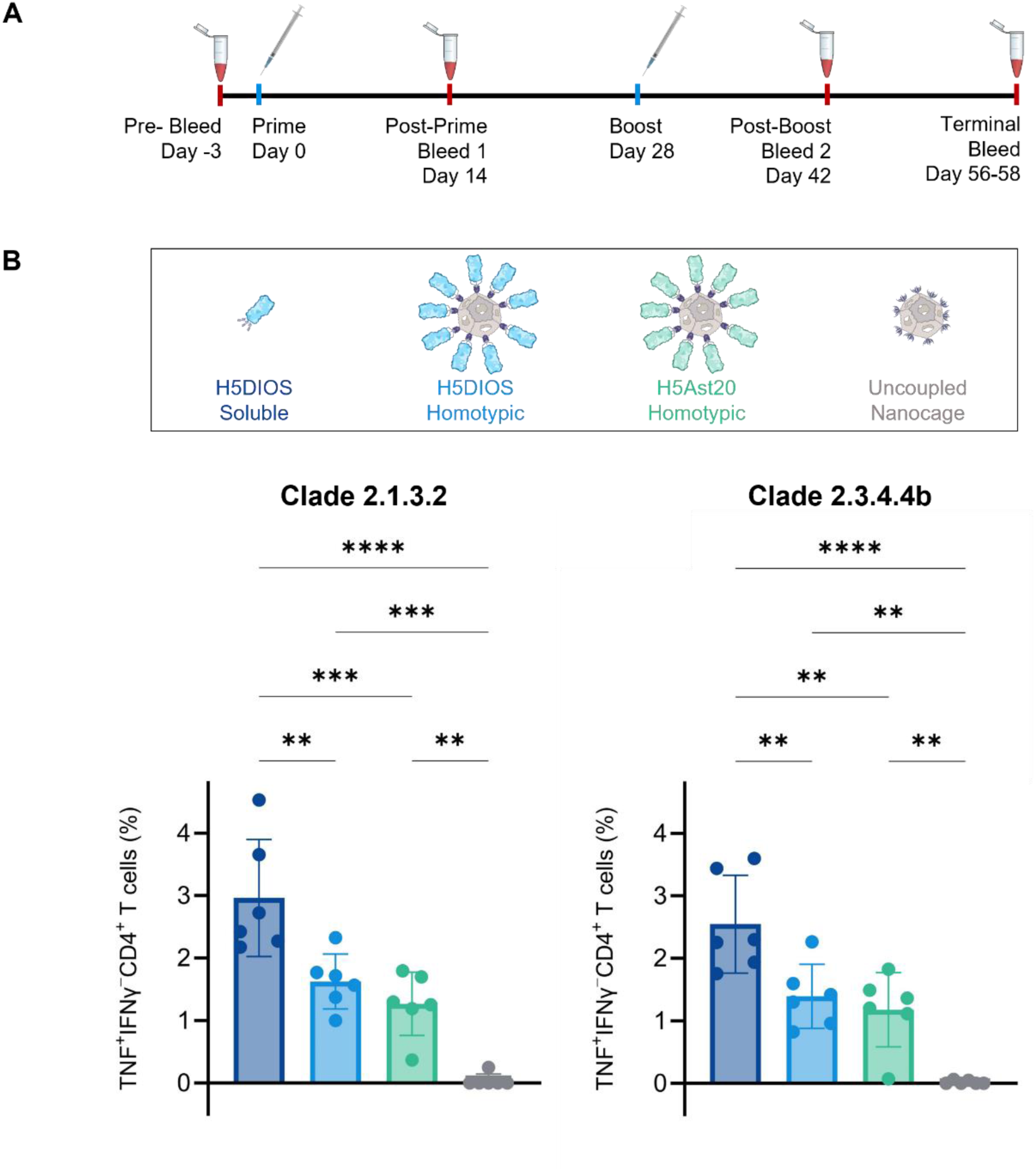
Broad CD4^+^ T cell response against antigenically distant H5 clades. **A)** Schedule of immunisation and blood sampling of mice. **B)** HPAI A/H5 HA-specific T cell responses quantified by TNF^+^IFNγ^−^CD4^+^ T cells. Splenocytes were collected from mice immunised with DIOSvax-H5_inter_ Soluble (dark blue), DIOSvax-H5_inter_ Homotypic Nanocage (light blue), H5Ast20 Homotypic Nanocage (green) or Uncoupled Nanocage (grey). Splenocytes were stimulated *ex vivo* with peptide pools for the HA of either the A/Indonesia/CDC835/2006 or the A/Aves/Guanajuato/CPA-18539-23/2023 strain. Cytokine production was measured by intracellular cytokine staining and flow cytometry. Each dot represents one mouse. The mean is denoted by a bar ± 1 s.d.; *n* = 6. Statistical significance was calculated by ANOVA, followed by Tukey’s multiple comparison post hoc test of % gated cells and plotted as \**P* < 0.05, \*\**P* < 0.01, \*\*\**P* < 0.001, \*\*\*\**P* < 0.0001.

Cross-reactive T cell responses targeting multiple conserved viral proteins have been shown to play important roles in mitigating influenza infection-associated pathology and protecting against severe disease^37,38^. Typically, CD8^+^ T cells are responsible for direct killing of infected cells, while CD4^+^ T cells predominantly augment CD8^+^ T cell functions and antibody responses during viral infections^37,38^. Interestingly, influenza-specific CD4^+^ cells have also been shown to mediate viral clearance^37,38^. HPAI A/H5 HA-specific CD4^+^ and CD8^+^ T cells were measured by intracellular cytokine staining for tumour necrosis factor (TNF) and interferon gamma (IFNγ), following *ex vivo* stimulation with commercially-available peptide pools of the clade 2.1.3.2 A/Indonesia/CDC835/2006 and the clade 2.3.4.4b A/Aves/Guanajuato/CPA-18539-23/2023 strains (Fig. 3b). DIOSvax-H5_inter_ Soluble antigen and the DIOSvax-H5_inter_ Homotypic Nanocages and H5Ast20 Homotypic Nanocages all induced significant TNF^+^IFNγ^−^CD4^+^ cells, compared to Uncoupled Nanocages upon stimulation with either peptide pool. Interestingly, mice immunised with DIOSvax-H5_inter_ Soluble antigens induced significantly higher percentages of TNF^+^IFNγ^−^CD4^+^ than the other two vaccine groups, indicating superior performance as a T cell immunogen. Antigen-specific T cells were not induced above background levels for all other assessed phenotypes (Fig. S3). TNF has been associated with potent anti-influenza activity, including stronger inhibition of viral replication than IFNγ^39^ and stimulation of cytokine and chemokine production by lung epithelial cells^40^. Thus, the vaccine-induced cytokines could play functionally important roles as a critical part of the host responses that contribute to overall protection.

### Computationally-designed HA nanocages neutralise multiple diverse H5 clades

Neutralising antibodies that block virus entry into host cells are well established as the primary immune correlate of protection for influenza vaccine effectiveness^41^. In recent years, pseudotype-based neutralisation assays have become widely accepted as a reliable surrogate for microneutralisation assays using live viruses^32,42^. We tested neutralisation of boosted sera at terminal bleed against a panel of H5 lentiviral pseudotypes^32,42^ from the 12 selected clades and subclades (Fig. 4). The H5Ast20 Homotypic Nanocage raised a potently neutralising immune response against the pseudotype from the circulating clade 2.3.4.4b. In addition to this strong matched response, the H5Ast20 Homotypic Nanocage induced similar levels of neutralisation against the closely related 2.3.4.4a and 2.3.4.4c strains. However, the H5Ast20 Homotypic Nanocage induced limited immune breadth and failed to induce significant neutralisation against several of the other tested pseudotypes when compared to Uncoupled Nanocage control (Fig. 4 and compiled in Fig. 5).

**Fig. 4.**
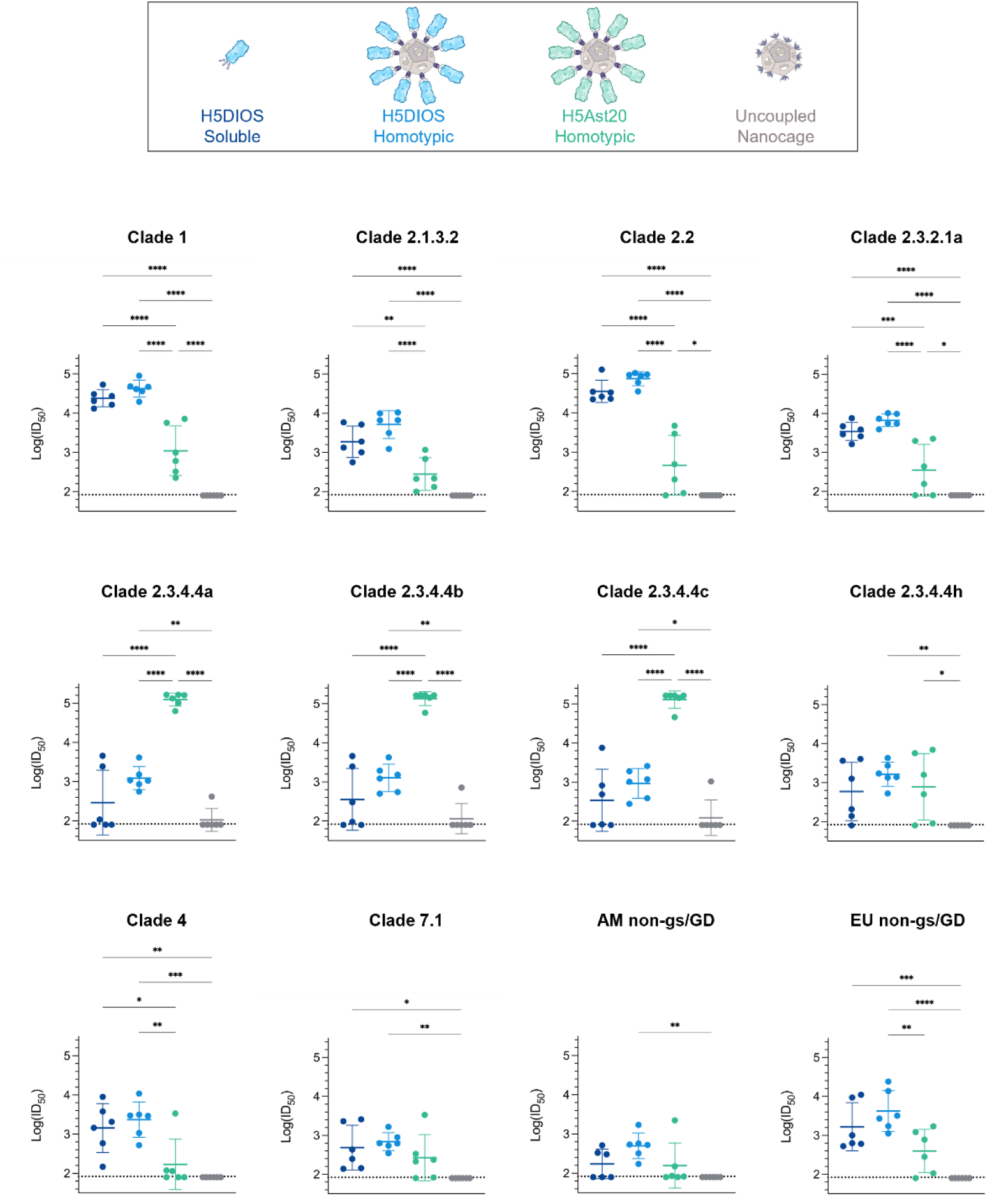
Broad neutralising antibody response against diverse H5 clades. Neutralisation of influenza pseudotypes by boosted mouse sera. Clade 1, 2.1.3.2, 2.2, 2.3.2.1a, 2.3.4.4a, 2.3.4.4b, 2.3.4.4c, 2.3.4.4h, 4, 7.1, AM non-gs/GD and EU non-gs/GD pseudotypes were tested for neutralisation by terminal bleed sera from mice immunised with DIOSvax-H5_inter_ (H5DIOS) Soluble (dark blue), DIOSvax-H5_inter_ Homotypic Nanocage (light blue), H5Ast20 Homotypic Nanocage (green) or Uncoupled Nanocage (grey). Each dot represents one mouse, showing the serum dilution giving 50% inhibition of infection (ID50). Dashed horizontal lines represent the limit of detection of log_10_(ID_50_) = 1.9. The mean is denoted by a horizontal line ± 1 s.d.; *n* = 6. Statistical significance was calculated by ANOVA, followed by Tukey’s multiple comparison post hoc test of log_10_(ID_50_) values and plotted as \**P* < 0.05, \*\**P* < 0.01, \*\*\**P* < 0.001, \*\*\*\**P* < 0.0001.

**Fig. 5.**
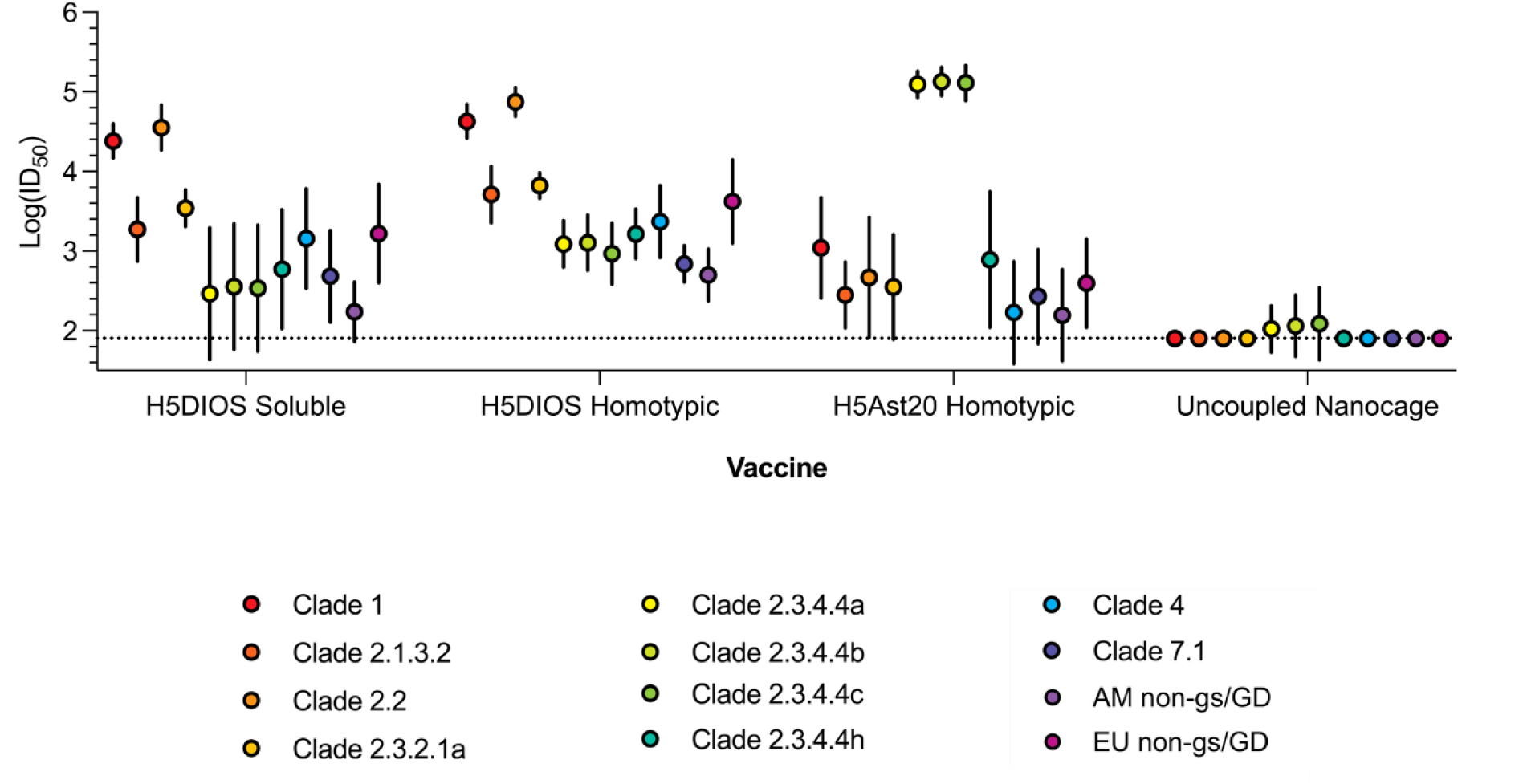
Further comparison of H5 neutralisation. Neutralisation of influenza pseudotypes by boosted mouse sera grouped by immunogen. Mice were immunised with DIOSvax-H5_inter_ (H5DIOS) Soluble, DIOSvax-H5_inter_ Homotypic Nanocage, H5Ast20 Homotypic Nanocage or Uncoupled Nanocage. Clade 1 (red), 2.1.3.2 (orange-red), 2.2 (orange), 2.3.2.1a (orange-yellow), 2.3.4.4a (yellow), 2.3.4.4b (yellow-green), 2.3.4.4c (green), 2.3.4.4h (aquamarine), 4 (light blue), 7.1 (dark blue), AM non-gs/GD (purple) and EU non-gs/GD (magenta) pseudotypes. The mean is shown ±1 s.d.; *n* = 6. Statistical significance is presented in Figure 4.

In contrast, the DIOSvax-H5_inter_ Homotypic Nanocage induced significant neutralisation against all of the tested pseudotypes. Every mouse immunised with the DIOSvax-H5_inter_ Homotypic Nanocage demonstrated neutralisation of every tested H5 pseudotype (Fig. 4). The DIOSvax-H5_inter_ Soluble antigen elicited high neutralisation comparable to the DIOSvax-H5_inter_ Homotypic Nanocage against HA pseudotypes from clade 1, 2.1.3.2, 2.2, 2.3.2.1a and the LPAI lineage EU non-gs/GD, but failed to consistently neutralise the currently circulating 2.3.4.4b and its closely related 2.3.4.4a, 2.3.4.4c and 2.3.4.4h strains (Fig. 4). Our results show that the DIOSvax-H5_inter_ Homotypic Nanocages, developed by combining two synthetic vaccine technologies, induced neutralising antibody responses with significantly greater breadth than either the multivalent display (H5Ast20 Homotypic Nanocage), or the computational design (DIOSvax-H5_inter_ Soluble) alone (Fig. 5). Potent neutralisation by DIOSvax-H5_inter_ Homotypic Nanocages was observed across all three main lineages of influenza HPAI and LPAI A/H5 viruses, representing unprecedented pan-H5 reactivity against diverse past and future strains.

## DISCUSSION

The ongoing panzootic of clade 2.3.4.4b HPAI A/H5 viruses has propelled global efforts in pre-pandemic vaccine design and stockpiling^43^. Historically stockpiled, U.S.-licensed inactivated HPAI A/H5 vaccines derived from the clade 1 A/Vietnam/1203/2004 and the clade 2.1.3.2 A/Indonesia/5/2005 strains have been tested for cross-reactivity against H5Ast20. Vaccinee sera from previous trials showed that cross-protective haemagglutination inhibition titres of ≥1:40 were only reached in 25-64% of individuals after multiple adjuvanted high-dose immunisations, highlighting the need for an update in stockpile to match the latest CVV^6^. In a recent study, the clade 2.3.4.4b inactivated and recombinant protein vaccines conferred 100% protection against lethal challenge by H5Ast20 in mice^7^. High-dose mRNA immunisation regimens have also demonstrated success in conferring homologous protection^8,9^. A nucleoside-modified mRNA vaccine for the same virus strain induced robust antibody and CD8^+^ T cell responses in mice and protected ferrets in a challenge study. Although cross-clade binding antibodies were detected in mice, the authors anticipated that they were likely targeting non-neutralising epitopes in the HA stalk and unlikely to provide sterilising immunity^8^. Another mRNA vaccine encoding the clade 2.3.4.4b A/chicken/Ghana/AVL-76321VIR7050-39/2021 strain cross-protected mice from heterologous challenge by the clade 2.3.2.1a A/India/SARI-4571/2021 strain. However, no neutralising antibody was detected for both the 1 μg and 10 μg dose groups against homologous and heterologous viruses, a key correlate of protection against infection^9^. Self-amplifying mRNA vaccines encoding full-length and truncated HAs of the clade 2.3.4 A/Anhui/1/2005 strain also induced binding antibodies against the matched strain but breadth and neutralisation were not evaluated^10^. Although multivalent mRNA-lipid nanoparticles designed as universal influenza priming vaccines are under clinical development, they are not expected to induce protective levels of neutralising antibody responses against heterologous viruses^44^. Even with the streamlined manufacturing process of mRNA vaccines, these strain-specific approaches are only applicable post hoc upon the precise characterisation of a pandemic strain. Hence, vaccines based on virus isolates of the past remain vulnerable to rapid and inevitable antigenic mismatches due to the highly mutagenic nature of influenza viruses.

To mitigate the uncertainty of zoonotic spillover in the pre-pandemic phase, next-generation vaccine, technologies have been explored for developing “future-proofed” immunogens with broader protective profiles. Antigenic cartography analysis of the eight subclades 2.3.4.4a-h identified a recombinant strain denoted H5-Re11_Q115L/R120S/A156T that was antigenically representative at clade-level only^45^. Multi-clade protection against matched virus challenge has also been achieved in chickens with a virus-like particle vaccine expressing the clade 2.3.2.1c and 2.3.4.4c HA from a retroviral gag protein backbone. However, no evidence of broader protection beyond the tested clades was reported^46^. More recently, a parainfluenza virus 5-based H5Ast20 vaccine induced sterilising immunity against heterologous clade 2.2, 2.3.2, 2.3.4 and 2.3.4.4h viruses in mice and ferrets^34^. Another viral-vectored vaccine used the Herpesvirus of Turkeys to express a recombinant H5 consensus representing multiple virus strains of the clade 2.2, which conferred full protection against challenge by mismatched clade 1 viruses^35^. While these studies have demonstrated limited breadth of protection against various clades within the HPAI lineage, more distantly related 4 and 7.1 clades and the LPAI lineages were not investigated.

Here, we present a combined novel approach using both computational antigen design technology, together with a multivalent nanocage delivery platform, both of which individually have elicited consistent, broadly protective responses against different rapidly evolving RNA viruses in multiple animal models^13,14,12,23,26–28^. A computational coronavirus antigen developed using the DIOSynVax technology^13^ and a human cytomegalovirus vaccine using the SpyTag/SpyCatcher technology have demonstrated adequate safety in rabbit toxicology screening and are currently in clinical trials in the UK^47^. Building on the success of these validated approaches, we have produced a dose-sparing and scalable *in silico* optimised DIOSvax-H5_inter_ Homotypic Nanocage vaccine candidate of paramount clinical relevance. Ongoing vaccine development efforts against clade 2.3.4.4b HPAI A/H5 viruses remain vulnerable to strain mismatching and immune evasion, especially in light of other HPAI A/H5 clades such as 2.3.2.1c and H5Nx reassortants that have infected humans in 2024^48^. By inducing robust pan-H5 humoral and cellular immunogenicity against a diverse range of H5 influenza clades, this vaccine strategy offers a proactive, robust response to currently circulating HPAI A/H5 viruses and to future evolutionarily-related strains with pandemic potential.

## MATERIALS AND METHODS

### Phylogenetic analysis

Influenza HA nucleotide sequences were downloaded from the GISAID database^19^ and employed for multiple sequence alignment using MUSCLE^49^. This alignment then provided the input for phylogenetic tree reconstruction using IQ-TREE^50^. The phylogenetic tree was then analysed with HyPhy^20^, allowing reconstruction of the DIOSvax-H5_inter_ design.

### Plasmids and cloning

Cloning was performed via standard PCR methods using the Q5 High-Fidelity PCR kit (New England Biolabs), Gibson assembly using the Gibson Assembly® Master Mix (New England Biolabs), and site-directed mutagenesis using the Q5® Site-Directed Mutagenesis Kit (New England Biolabs). The genes of interest were validated by Sanger sequencing (Source Bioscience).

pET28a-SpyCatcher003-mi3 (GenBank MT945417, Addgene 159995) was previously described^23^. pEVAC-H5Ast20-SpyTag003-His_6_ (GenBank deposition in progress) was created by inserting Influenza A/Astrakhan/3212/2020 H5 ectodomain (GISAID EPI1846961, residues 1-521), GSGSGSPGS linker, T4 bacteriophage fibritin foldon domain (GYIPEAPRDGQAYVRKDGEWVLLSTFL)^31^, (GGS)_3_G linker, SpyTag003 and His_6_ tag into the pEVAC plasmid^32^, followed by deletion of residues 341-344 (KRRK) to remove the polybasic cleavage site. pEVAC-DIOSvax-H5_inter_-SpyTag003-His_6_ was created by replacing H5Ast20 ectodomain with DIOSvax-H5_inter_ ectodomain (residues 1-522), followed by deletion of amino acids 341-345 (RRRKK) to remove the polybasic cleavage site.

### Bacterial protein expression of SpyCatcher003-mi3

Bacterial protein expression of the SpyCatcher003-mi3 nanocage was previously described^28^. *E. coli* BL21(DE3) cells (Agilent) were transformed with pET28a-SpyCatcher003-mi3 and were grown on LB-Agar plates containing 50 μg/mL kanamycin for 16 h at 37 °C. One colony from this plate was inoculated in 10 mL LB containing 50 μg/mL kanamycin and grown for 16 h at 37 °C with shaking at 200 rpm. This starter culture was added to 1 L LB containing 50 μg/mL kanamycin and incubated at 37 °C with shaking at 200 rpm until optical density (OD)_600_ was 0.6. Cultures were induced with 0.5 mM isopropyl β-D-1-thiogalactopyranoside and were grown for 16 h at 22 °C with shaking at 200 rpm, before centrifugation for 30 min at 4,000 g and 4 °C.

### Purification of SpyCatcher003-mi3

Purification of the SpyCatcher003-mi3 nanocage was previously described^28^. Cell pellets were resuspended in 20 mL 20 mM Tris-HCl, 300 mM NaCl, pH 8.5, that was supplemented with 0.1 mg/mL lysozyme, 1 mg/mL cOmplete mini EDTA-free protease inhibitor (Roche) and 1 mM phenylmethanesulfonyl fluoride (PMSF). The lysate was incubated for 45 min at 4 °C with end-over-end mixing. An Ultrasonic Processor equipped with a microtip (Cole-Parmer) was used for sonication on ice (four times for 60 s, 50% duty-cycle). Centrifugation was performed for 45 min at 35,000 g and 4 °C to clear cell debris. 170 mg of ammonium sulfate was added for every mL of lysate. The mixture was incubated for 1 h at 4 °C, with mixing at 120 rpm to precipitate the particles. Centrifugation for 30 min at 30,000 g and 4 °C was performed. The pellet was resuspended in 10 mL mi3 buffer (25 mM Tris–HCl, 150 mM NaCl, pH 8.0) at 4 °C. The solution was filtered sequentially through 0.45 μm and 0.22 μm syringe filters (Starlab). The filtrate was dialysed for 16 h against 1,000-fold excess of mi3 buffer. The dialysed particles were centrifuged for 30 min at 17,000 g and 4 °C and subsequently filtered through a 0.22 μm syringe filter. The purified SpyCatcher003-mi3 was loaded onto a HiPrep Sephacryl S-400 HR 16-600 column (GE Healthcare), which had been equilibrated with mi3 buffer using an ÄKTA Pure 25 system (GE Healthcare). The proteins were separated with collections at each 1 mL elution fraction. The fractions containing the purified particles were pooled and concentrated using a Vivaspin 20 100 kDa molecular weight cut-off (MWCO) centrifugal concentrator (GE Healthcare). The final nanocage was stored at −80 °C.

### Mammalian protein expression of HA-SpyTag003

Mammalian expression of all HA constructs was performed in Expi293F^TM^ cells (Thermo Fisher). Cells were cultured in Expi293^TM^ Expression Medium (Thermo Fisher) in suspension at 37 °C, 8% (v/v) CO_2_, ≥80% relative humidity and rotating at 125 rpm. Transient transfections were performed using the ExpiFectamine^TM^ 293 Transfection Kit (Thermo Fisher). Cells at 3 × 10^6^ cells/mL were transfected with 1 μg of plasmid DNA and 2.7 μL of ExpiFectamine^TM^ 293 per mL of culture. ExpiFectamine^TM^ 293 Transfection Enhancers 1 and 2 were added 20 h post-transfection. Cell supernatants were harvested 5 days post-transfection by centrifuging for 10 min at 4,000 g and 4 °C and sterile filtering through a 0.45 μm filter followed by a 0.22 μm filter (Merck). cOmplete mini EDTA-free protease inhibitor cocktail was added to the supernatants at 1 mg/mL (Roche).

### Purification of HA-SpyTag003

HA-SpyTag003 proteins were purified by nickel-nitrilotriacetic acid (Ni-NTA) affinity chromatography. Ni-NTA agarose (Qiagen) was washed with 2 × 10 column volumes (CV) of Ni-NTA buffer (50 mM Tris-HCl, 300 mM NaCl, pH 7.8 at 4 °C). Mammalian cell supernatants were supplemented with 10× Ni-NTA buffer (500 mM Tris-HCl, 3 M NaCl, pH 7.8 at 4 °C) at 10% (v/v) and incubated with Ni-NTA affinity matrix for 1 h at 4 °C with rolling. The mixtures were added to an Econo-Pac Chromatography Column (Bio-Rad), passed through by gravity filtration, and washed with 2 × 10 CV of Ni-NTA wash buffer (10 mM imidazole in Ni-NTA buffer). Ni-NTA elution buffer (200 mM imidazole in Ni-NTA buffer) was added to the column, incubated for 5 min and passed through by gravity filtration to elute proteins. A total of 6 × 1 CV elutions was performed. Total eluates were dialysed through 3.5 kDa MWCO Spectra-Por^®^ Float-A-Lyzer^®^ G2 (Spectrum Labs) for 16 h against 1,000-fold excess phosphate buffered saline (PBS) (137 mM NaCl, 2.7 mM KCl, 10 mM Na_2_HPO_4_, 1.7 mM KH_2_PO_4_, pH 7.4). Dialysed eluates were concentrated in a 20 mL 10 kDa Pierce™ Protein Concentrator (Thermo Fisher) by centrifuging for 5 min at 4,000 g and 4 °C.

## SDS-PAGE

All fractions collected during transfection harvest and affinity chromatography (supernatant, flow-through, washes 1-2 and elutions 1-6) and purified proteins were analysed by SDS-PAGE under reducing conditions using the Criterion™ Cell electrophoresis system (Bio-Rad). 2× Laemmli Sample Buffer (Bio-Rad) was supplemented with 2-mercaptoethanol at 5% (v/v) and mixed with samples at 50% (v/v). Diluted samples were heated for 5 min at 95 °C and resolved in a 12.5% Criterion™ Precast Tris-HCl gel (Bio-Rad). Electrophoresis was performed in Tris/Glycine/SDS buffer (Bio-Rad) for 70 min at 180 V. Gels were stained for 1 h with InstantBlue^®^ Coomassie Stain (Abcam) and destained for 16 h in MilliQ water. Imaging was performed with ChemiDoc XRS+ imager and ImageLab software (Bio-Rad).

### PNGase F digestion

De-glycosylation of HA was performed using the PNGase F kit (New England Biolabs). 2 µM of HA was heated with 1 µL of 10× Glycoprotein Denaturing Buffer (0.5% (w/v) SDS, 40 mM dithiothreitol) for 10 min at 100 °C. The denatured protein was chilled on ice for 1 min and centrifuged for 10 s at 2,000 g. 2 µL of 10× GlycoBuffer 2 (50 mM sodium phosphate, pH 7.5 at 25 °C), 2 µL of 10% (v/v) NP-40, 6 µL of MilliQ water and 1 µL of PNGase F at 500,000 units/mL were added and incubated for 1 h at 37 °C, before resolving in a 12.5% (w/v) Criterion™ Precast Tris-HCl gel (Bio-Rad) with Coomassie staining.

### Endotoxin depletion and quantification

SpyCatcher003-mi3 and HA-SpyTag003 proteins were depleted of endotoxins using Triton X-114 phase separation^51^. Triton X-114 was mixed with the proteins at 1% (v/v). The mixtures were incubated for 5 min on ice, then incubated for 5 min at 37 °C, and centrifuged for 1 min at 16,000 g and 37 °C. The top phase was transferred to a fresh tube. A total of three repetitions were performed, followed by a final repetition without the addition of Triton X-114. All samples were assessed for final endotoxin concentrations using Pierce™ Chromogenic Endotoxin Quant Kit (Thermo Fisher) and confirmed to be below the accepted <20 Endotoxin Units (EU) per mL for vaccine products^52^.

### Immunogen preparation

The concentration of SpyCatcher003-mi3 nanocages was determined by A_280_ measurements taken using a NanoDrop™ One Spectrophotometer (Thermo Fisher). The concentration of HA-SpyTag003 was measured using the Pierce™ Bicinchoninic Acid Assay kit (Thermo Fisher). Doses were normalised by the number of antigens, to facilitate an equimolar amount of SpyCatcher003-mi3 nanocages with similar occupancy in each condition. 1.5 μM HA-SpyTag003 was incubated with 1 μM SpyCatcher003-mi3 for 24 h at 4 °C in PBS, pH

7.2. Uncoupled HA-SpyTag003 was incubated for 24 h at 4 °C in PBS, pH 7.2, without the addition of SpyCatcher003-mi3. Post-coupling, immunogens were analysed by SDS–PAGE/Coomassie, DLS and ELISA. Immunogens were further diluted in PBS, pH 7.2, to the final doses of 0.5 μg of total SpyTagged antigens, which relates to 8 pmol of uncoupled HA. Before immunisation, immunogens were mixed 1:1 with AddaVax (Invivogen).

### Dynamic light scattering (DLS)

Samples were centrifuged for 30 min at 16,000 g and 4 °C, before 70 μL was loaded onto a plastic cuvette. Samples were measured in triplicate at 20 °C using a Zetasizer Nano S (Malvern Panalytical) with 11 scans of 10 s each. Before collecting data, the cuvette was incubated in the instrument for 2 min to allow the sample temperature to stabilise. The size distribution was determined by the Zetasizer Software v.7.13 (Malvern Panalytical), calculating the mean and s.d. from the multiple scans.

### Transmission electron microscopy (TEM)

Samples were centrifuged for 30 min at 16,000 g at 4 °C. Carbon 400 mesh copper grids (EM Resolutions) were processed in a Quorum Emitech K100X glow discharger at 25 mA for 2 min. Samples were applied to the grids for 30 s, washed 2 × with distilled H_2_O for 10 s and stained with 2% (w/v) uranyl acetate for 10 s. Samples were blotted with filter paper between steps and air-dried after staining. Grids were imaged using a FEI Tecnai G2 80–200-keV transmission electron microscope at 200 keV with a 20 μm objective aperture at the Cambridge Advanced Imaging Centre.

## ELISA

Nunc™ MaxiSorp™ 96-Well flat-bottom plates (Thermo Fisher) were coated with 50 μL per well of 1 μg/mL coupled or uncoupled HA-SpyTag003 in PBS, pH 7.2, and incubated overnight at 4 °C. Plates were blocked with 3% (w/v) skimmed milk in PBS supplemented with 0.1% (v/v) Tween 20 (PBST) for 2 h at RT. Sera and monoclonal antibodies were serially diluted in 1% (w/v) skimmed milk in PBST using an eight-point, three-fold series starting at 1:200 and 0.5 μg/mL, respectively. After removing the blocking buffer, 50 μL per well of sera or broadly neutralising monoclonal antibodies CR9114^36^, were added and incubated for 2 h at RT with shaking at 350 rpm. Plates were washed 3 × with 200 μL per well of PBST. 50 μl of 1:3,000 (v/v) Peroxidase AffiniPure™ Goat Anti-Mouse IgG (H+L) (Jackson ImmunoResearch, 115-035-003) or 1:5,000 (v/v) Peroxidase AffiniPure™ Rabbit Anti-Human IgG (H+L) (Jackson ImmunoResearch, 309-035-003) per well was added and incubated for 1 h at RT with shaking at 350 rpm. Plates were washed 3 × with 200 μL per well of PBST. 50 μL per well of 1-Step Ultra TMB chromogenic substrate (Merck) was added to the plates incubated for 3 min at RT, before the chemical reaction was stopped with 50 μL 1 N H_2_SO_4_. OD_450_ was measured using BioTek 800 TS Absorbance Reader and Gen6 software (Agilent). Using GraphPad Prism (GraphPad Software v10.3.1), nonlinear regression was performed to plot sigmoidal dose-response curves, from which area under the curve (AUC) values were determined and plotted as means ± 1 s.d. of duplicate measurements.

### Mouse immunisation and blood sampling

Animal experiments were performed according to the UK Animals (Scientific Procedures) Act 1986, under Project Licence (PP9157246) and approved by the University of Cambridge Animal Welfare and Ethical Review Body. Fourteen groups of six 8–10-week-old female BALB/c mice were obtained from Charles River Laboratories. Mice were immunised twice at a 28-day interval. A total volume of 100 µL of 0.5 μg of HA-SpyTag003 was administered via the intramuscular route over both hind legs. Blood was sampled from the saphenous vein at 14-day intervals and animals were terminally bled by cardiac puncture under non-recovery anaesthesia on day 56. Whole blood was allowed to clot at 25 °C for 1-2 h and centrifuged for 5 min at 10,000 g. Clarified sera were transferred to fresh tubes, heat-inactivated for 30 min at 56 °C, and stored at −30 °C.

### Lentiviral pseudotype production

Influenza pseudotyped viruses A/Vietnam/1203/2004 (clade 1), A/Indonesia/5/2005 (clade 2.1.3.2), A/whooper swan/Mongolia/244/2005 (clade 2.2), A/Hubei/1/2010 (clade 2.3.2.1a), A/Sichuan/26221/2014 (clade 2.3.4.4a), A/Astrakhan/3212/2020 (clade 2.3.4.4b), A/gyrfalcon/Washington/41088-6/2014 (clade 2.3.4.4c), A/Guangdong/18SF020/2018 (clade 2.3.4.4h), A/goose/Guiyang/337/2006 (clade 4), A/chicken/Vietnam/NCVD-016/2008 (clade 7.1), A/chicken/Mexico/07/2007 (AM non-GS/GD) and A/mallard duck/Netherlands/41/2015 (EU non-GS/GD) were produced in HEK293T/17 cells as previously described^32,42^. Cells were cultured in DMEM GlutaMAX^TM^ (Thermo Fisher) supplemented with 10% (v/v) foetal bovine serum (FBS), 100 U/mL penicillin and 100 µg/mL streptomycin at 37 °C, 5% (v/v) CO_2_, ≥80% relative humidity. Transient transfections with lentiviral packaging plasmids p8.91-gag-pol^53^ and pCSFLW-firefly luciferase^54^, protease-bearing plasmid pCMV-TMPRSS4^55^ and glycoprotein-bearing plasmid pEVAC-HA were performed using the FuGENE^®^ HD Transfection Reagent (Promega). Cells at 4 × 10⁵ cells/mL were seeded in 6-well plates and incubated for 24 h at 37 °C until ∼70% confluence. Cells were transfected with 125 ng of p8.91, 187.5 ng of pCSFLW, 0-62.5 ng of pCMV-TMPRSS4 and 5-125 ng of pEVAC-HA plasmid DNA per mL of culture with 3 µL of FuGENE HD per µg of DNA. Exogenous neuraminidase from *Clostridium perfringens* (Merck) was added 24 h post-transfection at 0.5 U/ mL. Cell supernatants were harvested 48 h post-transfection by sterile filtering through a 0.45 μm filter (Merck) and stored at −80 °C. Pseudotypes were titrated on HEK293T/17 cells for 48 h at 37 °C using an eight-point, twofold series starting at 1:2 to measure relative luminescence units (RLU)/mL.

### Pseudotype-based microneutralisation assay

Pseudotype-based microneutralisation assays (pMN) were performed as previously described^32,42^. Sera and the broadly neutralising monoclonal antibody FI6^56^ were serially diluted in Nunc™ MicroWell™ 96-Well flat-bottom plates (Thermo Fisher) using a twelve-point, twofold series starting at 1:80 and 0.5 μg/mL, respectively. Influenza HA-bearing pseudotyped viruses at 1.5-5 × 10^6^ RLU/mL were added and incubated for 1 h at 37 °C and 5% (v/v) CO_2_. HEK293T/17 cells at 1.5 × 10^4^ cells/well were then added and incubated for 48 h at 37 °C and 5% (v/v) CO_2_. After removing the cell supernatants, excess Bright-Glo^TM^ (Promega) was added and incubated for 5 min at RT. Luminescence was measured using GloMax^®^ Explorer (Promega). Individual data points were normalised to 100% and 0% neutralisation controls, corresponding to values derived from uninfected cells and cells infected with pseudotyped virus in the absence of serum, respectively. Nonlinear regression analysis was performed to plot sigmoidal dose-response curves and determine half-maximal inhibitory dilution (ID_50_) values in log_10_ scale. Statistical significance of differences in log_10_(ID_50_) values between groups was determined using the ANOVA test, followed by the Tukey’s multiple comparison post hoc test. Results were plotted using GraphPad Prism (GraphPad Software v.10.3.1).

### Intracellular cytokine staining and flow cytometry

Single-cell suspension of murine splenocytes was prepared as previously described^57^. Peptide pools consisting of 15 amino acid peptides with 11 amino acid overlap spanning the H5N1 A/Indonesia/CDC835/2006 (clade 2.1.3.2; Swiss-Prot ID: A1BK62) and A/Aves/Guanajuato/CPA-18539-23/2023 (clade 2.3.4.4b; Swiss-Prot ID: WPD27583.1) HA proteins were used for stimulation (JPT peptides). Cells were treated by 50 U/mL of Benzonase**^®^**nuclease (Sigma Aldrich) in cell culture medium (RPMI 1640 Medium GlutaMAX™ (Thermo Fisher) supplemented with 1 mM sodium pyruvate, 50 µM 2-mercaptoethanol, 10% (v/v) FBS, 100 U/mL penicillin and 100 µg/ml streptomycin) to prevent clumping. Cells were resuspended in Nunc™ MicroWell™ 96-Well U-bottom plates (Thermo Fisher) at 5 × 10^5^ cells/well in cell culture medium. 2 µg/mL peptides or 1× eBioscience™ Cell Stimulation Cocktail (Thermo Fisher) were added and incubated for 2 h at 37 °C, 5% (v/v) CO_2_, ≥80% relative humidity. 1× eBioscience™ Protein Transport Inhibitor Cocktail (Thermo Fisher) was added and incubated for another 4 h. Plates were centrifuged for 3 min at 330 g to remove supernatant. Cells were washed with flow cytometry buffer (PBS supplemented with 2.5% (v/v) FBS and 2 mM EDTA). For surface marker staining, cells were incubated with 1:1,000 (v/v) eBioscience™ Fixable Viability Dye eFluor™ 780 (Thermo Fisher), 1:100 (v/v) CD3e Monoclonal Antibody (145-2C11) PE-Cyanine5.5 (Thermo Fisher, 35-0031-82), 1:100 (v/v) CD8a Monoclonal Antibody (53-6.7) PE-Cyanine7 (Thermo Fisher, 25-0081-82) and 1:100 (v/v) CD4 Monoclonal Antibody (RM4-5) APC (Thermo Fisher, 17-0042-82) in flow cytometry buffer for 30 min at 4°C. Cells were washed twice with flow cytometry buffer, fixed in Fixation/Permeabilization solution (BD Biosciences) for 10 min at RT and washed twice with flow cytometry buffer. For intracellular cytokine staining, cells were incubated with 1:100 (v/v) eBioscience™ IFN gamma Monoclonal Antibody (XMG1.2) Alexa Fluor™ 488 (Thermo Fisher, 53-7311-82) and 1:100 (v/v) eBioscience™ TNF alpha Monoclonal Antibody (MP6-XT22) PE-eFluor™ 610 (Thermo Fisher, 61-7321-82) in 1× Perm/Wash buffer (BD Biosciences) for 30 min at RT. Cells were washed twice with BD Perm/Wash™ buffer (BD Biosciences) and resuspended in flow cytometry buffer. Plates were analysed using the Attune™ Nxt Flow Cytometer (Thermo Fisher). Data analysis was performed in Attune Cytometric Software 6.21 (Thermo Fisher). Values less than zero after subtraction of background were assigned a value of zero. Statistical significance of differences in % gated cells between groups was determined using the ANOVA test, followed by the Tukey’s multiple comparison post hoc test. Results were plotted using GraphPad Prism (GraphPad Software v.10.3.1).

## ACKNOWLEDGEMENTS

We thank Dr. Katherine Stott from the University of Cambridge Department of Biochemistry Biophysical Suite for help with biophysical analysis, as well as Dr. Karin Mueller and Georgina Lindop from the Cambridge Advanced Imaging Centre for help with TEM. We thank Joey Olivier from the Laboratory of Viral Zoonotics of the University of Cambridge Department of Veterinary Medicine for help with splenocyte purification. C.Q.H. received funding from the Cambridge Commonwealth, European & International Trust, Croucher Foundation and St. John’s College, Cambridge. R.A.H. received funding from the Rhodes Trust, Townsend-Jeantet Prize Charitable Trust and St. John’s College, Oxford. M.R.H. and L.S.T. received funding from Flu Lab. J.L.H., G.W.C. and S.V. received funding to develop universal vaccine antigens from Flu Lab/Bill & Melinda Gates Foundation (INV-029365) and InnovateUK (DIOS-PIVa 105078) grants.

## AUTHOR INFORMATION

### Contributions

C.Q.H. conceived the main ideas of the study and acquired funding. C.Q.H. and R.A.H. designed and coordinated the project. L.S.T., M.R.H. and J.L.H. acquired funding and supervised the project. S.D.W.F. developed the computational pipeline and designed the antigen. J.L.H., S.V. and G.W.C. tested, validated and selected the candidate antigen breadth of immunogenicity by gene delivery. C.Q.H. and R.A.H. performed DNA cloning, protein production and purification, prepared immunogens, and completed SDS-PAGE, DLS, TEM and ELISA. C.Q.H., G.W.C., L.O. and P.T. performed mouse immunisation and blood sampling. C.Q.H. produced lentiviral pseudotypes with guidance from G.W.C. and N.T.. C.Q.H. and G.W.C. performed pseudotype-based microneutralisation. C.Q.H. and A.C.Y.C. performed splenocyte purification. C.Q.H. and E.T.A. performed intracellular cytokine staining and flow cytometry. S.V. and P.P. contributed to data analysis and visualisation. C.Q.H. wrote the manuscript. All authors contributed substantially to discussion of the content, reviewed and/or edited the manuscript before submission.

## ETHICAL DECLARATIONS

### Data availability

Further information and request for resources and reagents should be directed to and will be fulfilled by the lead contacts, J.L.H., M.R.H., and L.S.T.

### License information

For the purpose of Open Access, the author has applied a CC BY public copyright licence to any Author Accepted Manuscript (AAM) version arising from this submission.

