## Supplementary Information for "Digitally immune optimised haemagglutinin with nanocage plug-and-display elicits broadly neutralising pan-H5 influenza subtype vaccine responses"

#These authors contributed equally.

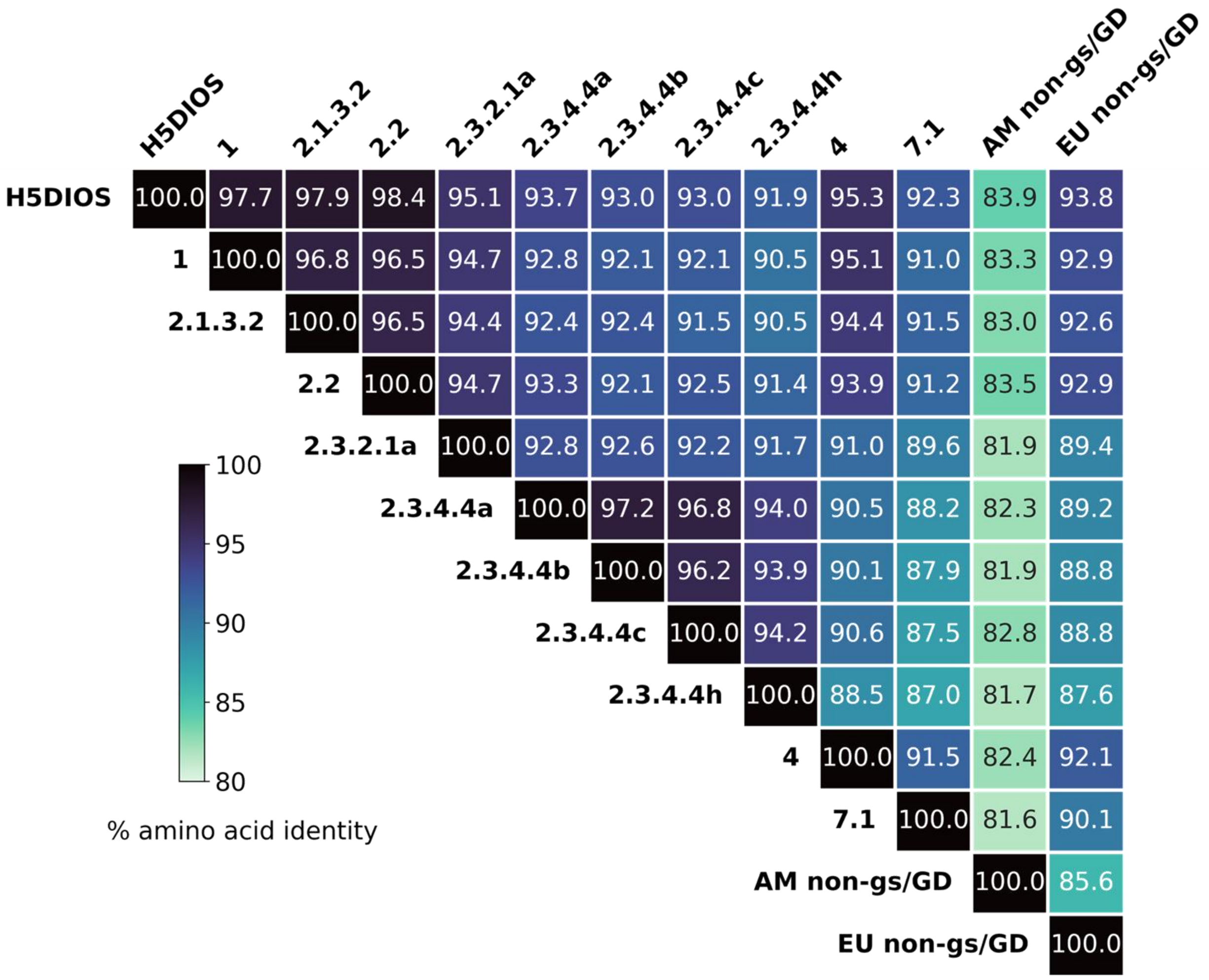


**Supplementary Figure 1. H5 Residue Conservation.** Heat map of percent HA amino acid sequence identity of H5 viruses used in this study.

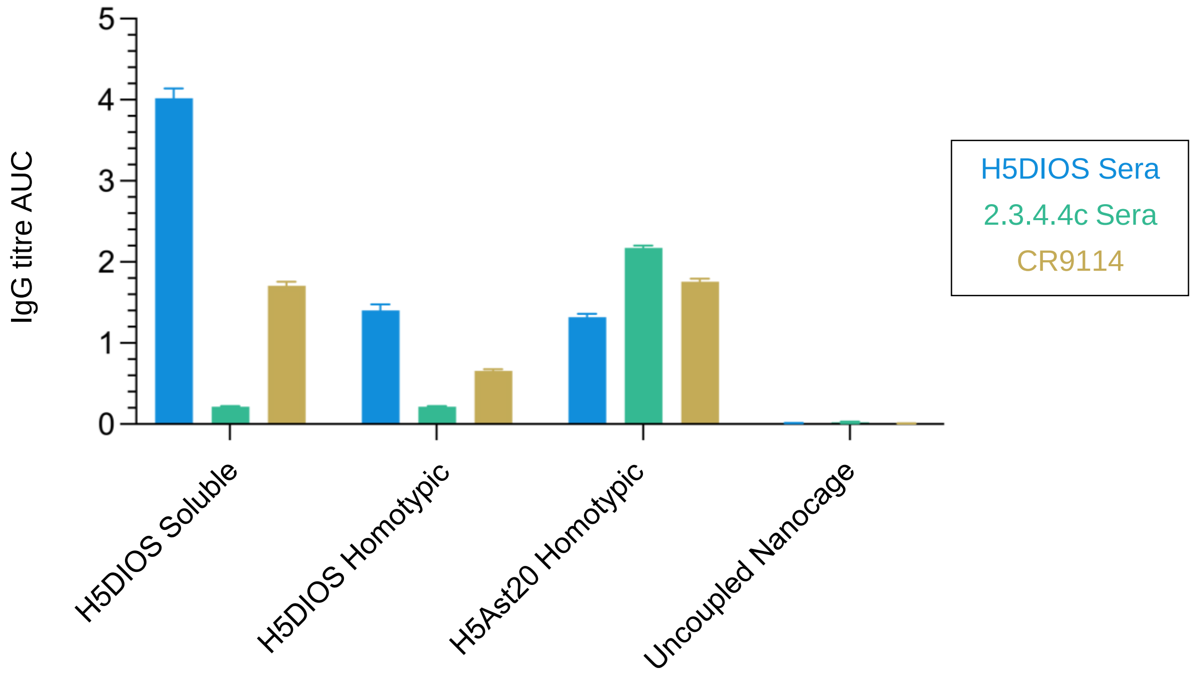


**Supplementary Figure 2. Validation of antigenicity of vaccine antigens by ELISA.** Binding of immobilised antigens to the broadly neutralising antibody CR9114 (yellow) and sera from mice previously immunized with DIOSvax-H5$_{inter}$ (H5DIOS) (blue) and 2.3.4.4c (green) as a DNA vaccine. The antigens tested were DIOSvax-H5$_{inter}$ Soluble, H5DIOS Homotypic Nanocage, H5Ast20 Homotypic Nanocage, and Uncoupled Nanocage. Results are represented as area under the curve (AUC) of a serial dilution. The mean is denoted by a horizontal line + 1 s.d.; $n = 2$.

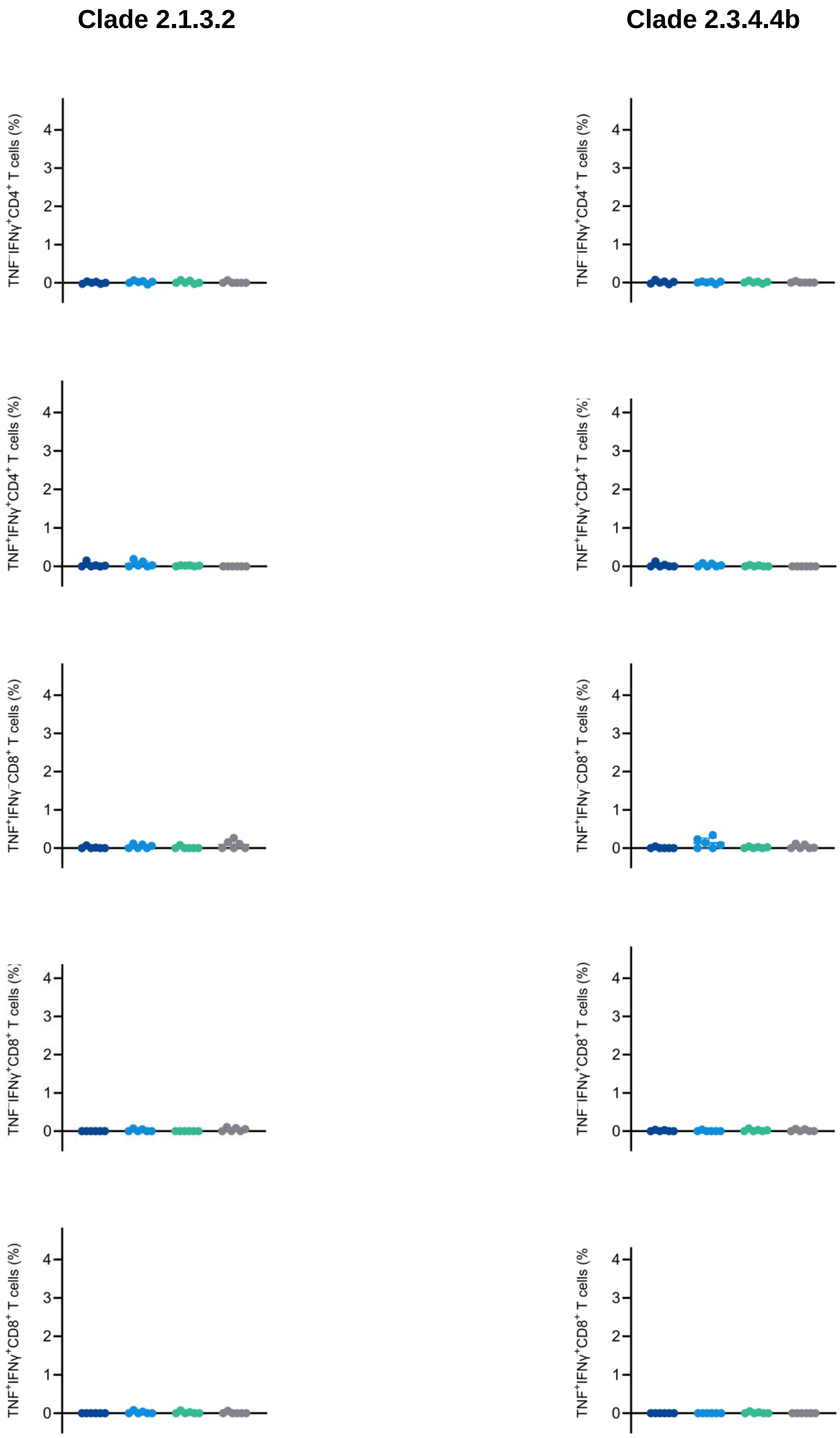


**Supplementary Figure 3.** Intracellular cytokine staining of HPAI A/H5 HA-specific T cells from mice immunised with DIOSvax-H5$_{inter}$ Soluble (dark blue), DIOSvax-H5$_{inter}$ Homotypic Nanocage (light blue), H5Ast20 Homotypic Nanocage (green) or Uncoupled Nanocage (grey). Splenocytes were stimulated *ex vivo* with peptide pools for the HA of either the A/Indonesia/CDC835/2006 or A/Aves/Guanajuato/CPA-18539-23/2023 strain. Each dot represents one mouse. The mean is denoted by a bar ± 1 s.d.; $n = 6$. Statistical significance was calculated by ANOVA, followed by Tukey's multiple comparison post hoc test of % gated cells. All comparisons were non-significant.
